# NudCL2 is an Hsp90 cochaperone to regulate sister chromatid cohesion by stabilizing cohesin subunits

**DOI:** 10.1101/358911

**Authors:** Yuehong Yang, Wei Wang, Min Li, Wen Zhang, Yuliang Huang, Ya Gao, Wei Zhuo, Xiaoyi Yan, Wei Liu, Fangwei Wang, Dingwei Chen, Tianhua Zhou

## Abstract

Sister chromatid cohesion plays a key role in ensuring precise chromosome segregation during mitosis, which is mediated by the multisubunit complex cohesin. However, the molecular regulation of cohesin subunits stability remains unclear. Here, we show that NudCL2 (NudC-like protein 2) is essential for the stability of cohesin subunits by regulating Hsp90 ATPase activity in mammalian cells. Depletion of NudCL2 induces mitotic defects and premature sister chromatid separation and destabilizes cohesin subunits that interact with NudCL2. Similar defects are also observed upon inhibition of Hsp90 ATPase activity. Interestingly, ectopic expression of Hsp90 efficiently rescues the protein instability and functional deficiency of cohesin induced by NudCL2 depletion, but not *vice versa*. Moreover, NudCL2 not only binds to Hsp90, but also significantly modulates Hsp90 ATPase activity and promotes the chaperone function of Hsp90. Taken together, these data suggest that NudCL2 is a previously undescribed Hsp90 cochaperone to modulate sister chromatid cohesion by stabilizing cohesin subunits, providing a hitherto unrecognized mechanism that is crucial for faithful chromosome segregation during mitosis.

## Introduction

During mitosis, the proper segregation of duplicated chromosomes into daughter cells is required for maintaining genome integrity. Sister chromatid cohesion, which is mediated by the highly conserved protein complex cohesin, plays an essential role in chromosome segregation [1, 2]. In general, the canonical cohesin complex is composed of four subunits, including two structural maintenance of chromosomes proteins (SMCs) and two sister chromatid cohesion proteins (SCCs). Two large ATPases (SMC1 and SMC3) are joined together by th e α-kleisin subunit Scc1/Mcd1/Rad21 to form a ring-shaped structure that mediates the cohesion by embracing sister chromatids. Stromal antigen (SA), the Scc3 homologue in vertebrate somatic cells, directly interacts with Scc1 via its C-terminal region to stabilize the cohesin ring.

The cohesin complex is dynamically regulated by various factors during cell cycle progression in vertebrate cells [1]. In telophase and G1, cohesin is loaded onto chromatin by the Scc2-Scc4 complex, which is counteracted by Wapl (wings-apart like) [3, 4]. During S phase, SMC3 is acetylated by Eco1 (establishment of cohesion protein 1) to enhance the cohesin-DNA interaction and promote the recruitment of sororin to cohesin. This recruitment displaces Wapl from its binding partner Pds5 (precocious dissociation of sisters protein 5) to further stabilize the cohesin ring that entraps the sister chromatids during S/G2 phase [5, 6]. In early mitosis, the phosphorylation of SA2 by polo-like kinase 1 and the opening of the cohesin ring by Wap1 result in the removal of the majority of cohesin from the chromosome arms [7, 8]. At the same time, the centromeric cohesin is protected by the Sgo1-PP2A (shugoshin 1 and protein phosphatase 2A) complex and Haspin (histone H3 associated protein kinase) until metaphase [9–14]. The cleavage of Scc1 by separase 4 and the deacetylation of SMC3 by histone deacetylase HOS1 at the metaphase-anaphase transition promote the complete removal of cohesin from chromatin [15–19]. This precise regulation of the cohesin complex ensures the accurate chromosome segregation in cell cycle progression. However, little is known about the regulation of cohesin subunits stability.

NudC (nuclear distribution gene C) is a protein that is highly conserved from yeasts to humans [20–22]. In the filamentous fungus *Aspergillus nidulans*, NudC has been demonstrated as an upstream regulator of NudF (a homologue of human LIS1, a key regulator of cytoplasmic dynein) [20, 23, 24]. We cloned and characterized two mammalian homologues of NudC, NudCL (NudC-like protein) and NudCL2 [25, 26]. NudC and NudCL participate in many biological processes, including cell cycle progression and cell migration [25, 27–30]. Recently, our group found that NudCL2 is able to stabilize LIS1 by enhancing the interaction between LIS1 and Hsp90 [26]. All of the NudC homologues share a conserved core domain of p23 (p23 domain), which modulates Hsp90 ATPase activity to promote the folding of client proteins, and are involved in the regulation of protein stability [25, 26, 28, 31–33]. However, whether the members of NudC family are the Hsp90 cochaperones is largely unknown.

The cohesin complex is dynamically regulated by various factors during cell cycle progression in vertebrate cells [1]. In telophase and G1, cohesin is loaded onto chromatin by the Scc2-Scc4 complex, which is counteracted by Wapl (wings-apart like) [3, 4]. During S phase, SMC3 is acetylated by Eco1 (establishment of cohesion protein 1) to enhance the cohesin-DNA interaction and promote the recruitment of sororin to cohesin. This recruitment displaces Wapl from its binding partner Pds5 (precocious dissociation of sisters protein 5) to further stabilize the cohesin ring that entraps the sister chromatids during S/G2 phase [5, 6]. In early mitosis, the phosphorylation of SA2 by polo-like kinase 1 and the opening of the cohesin ring by Wap1 result in the removal of the majority of cohesin from the chromosome arms [7, 8]. At the same time, the centromeric cohesin is protected by the Sgo1-PP2A (shugoshin 1 and protein phosphatase 2A) complex and Haspin (histone H3 associated protein kinase) until metaphase [9–14]. The cleavage of Scc1 by separase and the deacetylation of SMC3 by histone deacetylase HOS1 at the metaphase-anaphase transition promote the complete removal of cohesin from chromatin [15–19]. This precise regulation of the cohesin complex ensures the accurate chromosome segregation in cell cycle progression. However, little is known about the regulation of cohesin subunits stability.

NudC (nuclear distribution gene C) is a protein that is highly conserved from yeasts to humans [20–22]. In the filamentous fungus *Aspergillus nidulans*, NudC has been demonstrated as an upstream regulator of NudF (a homologue of human LIS1, a key regulator of cytoplasmic dynein) [20, 23, 24]. We cloned and characterized two mammalian homologues of NudC, NudCL (NudC-like protein) and NudCL2 [25, 26]. NudC and NudCL participate in many biological processes, including cell cycle progression and cell migration [25, 27–30]. Recently, our group found that NudCL2 is able to stabilize LIS1 by enhancing the interaction between LIS1 and Hsp90 [26]. All of the NudC homologues share a conserved core domain of p23 (p23 domain), which modulates Hsp90 ATPase activity to promote the folding of client proteins, and are involved in the regulation of protein stability [25, 26, 28, 31–33]. However, whether the members of NudC family are the Hsp90 cochaperones is largely unknown.

In this study, we provide evidence that NudCL2 is required for stabilizing cohesin subunits and ensuring accurate sister chromatid separation during mitosis. Our data reveal that NudCL2 significantly inhibits Hsp90 ATPase activity and enhances its chaperone function. Moreover, ectopic expression of Hsp90 efficiently rescues the cohesion defects in NudCL2-depleted cells. Thus, we propose that NudCL2 acts as an Hsp90 cochaperone to maintain cohesin subunit stability, which is crucial for faithful chromosome segregation in mammalian cells.

## Results

### Depletion of NudCL2 induces chromosome misalignment

To address the role of NudCL2 in mitosis, we employed small interfering RNAs (siRNAs) to deplete NudCL2 in HeLa cells. We used two siRNA oligos targeting two different regions of *NudC*L2 mRNA (NudCL2 siRNA and NudCL2 siRNA-2). We found that the protein level of NudCL2 was substantially reduced 72 h post-transfection (Fig 1A and Appendix Fig S1A). Immunofluorescence microscopy showed that downregulation of NudCL2 led to the accumulation of mitotic cells (Fig 1B and C; Appendix Fig S1B and C). Further analysis revealed that the percentage of mitotic cells with misaligned chromosomes was significantly higher in NudCL2-depleted cells than in control cells (Fig 1D and Appendix Fig S1D). Importantly, these defects in cells depleted of NudCL2 were effectively rescued by ectopic expression of siRNA-resistant NudCL2 (Appendix Fig S2). In addition, knockdown of NudCL2 in nontumor HEK-293 cells also resulted in mitotic arrest and chromosome misalignment (Appendix Fig S3). Collectively, these data suggest that NudCL2 plays an important role in mitosis.

**Figure 1.**
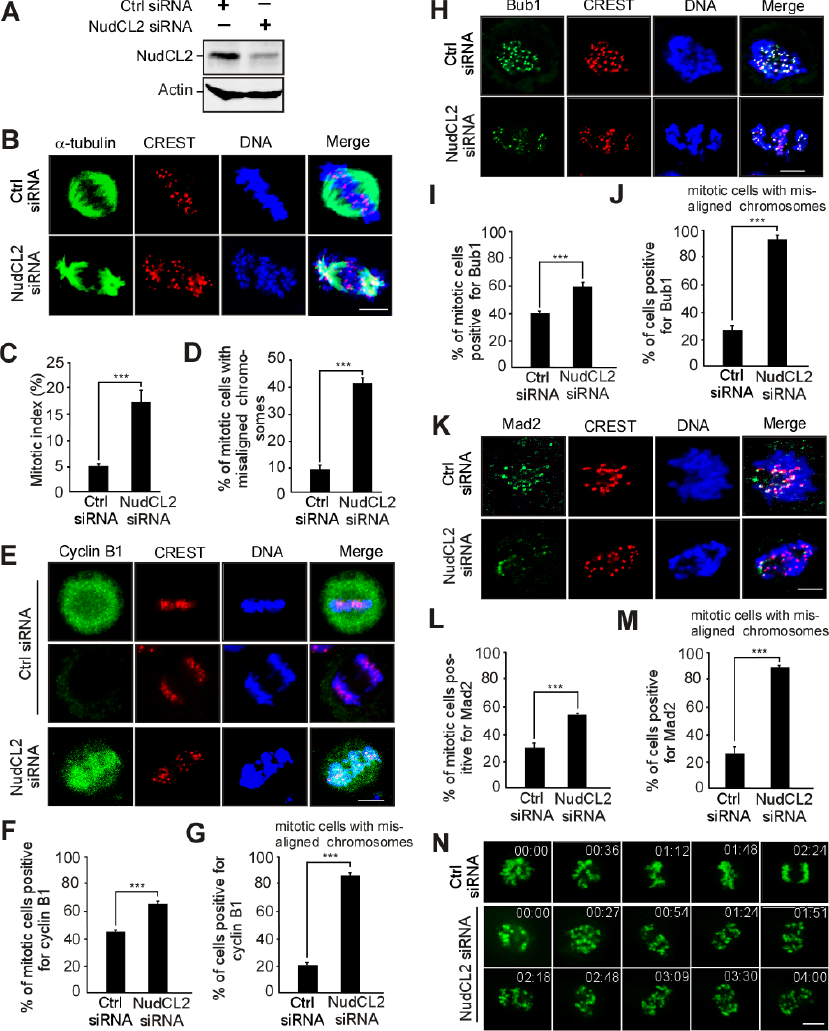
Depletion of NudCL2 causes mitotic defects. HeLa cells were transfected with either control or NudCL2 siRNA for 72 h and subjected to the following analyses. A. Western blot analysis showed efficient suppression of NudCL2. Actin was used as a loading control. B-D Immunofluorescence analysis displayed misalignment of chromosomes in NudCL2-depleted cells. Cells were stained with an anti-α-tubulin antibody and human CREST serum (B). The mitotic index was calculated (C). The percentage of mitotic cells with misaligned chromosomes was significantly higher in NudCL2-depleted cells than in control cells (D). E-G Immunostaining revealed that NudCL2-depleted cells exhibiting misaligned chromosomes were positive for cyclin B1 staining. Cells were probed with an anti-cyclin B1 antibody and human CREST serum (E). Mitotic cells positive for cyclin B1 were calculated (F). The frequency of cyclin B1-positive mitotic cells with misaligned chromosomes was plotted (G). H-M The spindle checkpoint proteins Bub1 and Mad2 were present at the kinetochores of NudCL2-depleted cells with misaligned chromosomes. Cells were stained with the indicated antibodies and human CREST serum. Mitotic cells positive for Bub1 (I) or Mad2 (L) were calculated. Mitotic cells with misaligned chromosomes with Bub1 (J) or Mad2 (M) signals were also counted. N Live imaging of HeLa cells stably expressing GFP-H2B showed that some chromosomes failed to congress to the metaphase plate in NudCL2-depleted cells. Time is shown as hh:mm. Data information: DNA was visualized with DAPI. Scale bars, 10 μm. Quantitative data are expressed as the mean ± SD (at least three independent experiments). More than 180 cells were counted in each experiment. \*\*\**p* < 0.001, Student’s *t* test.

To determine whether the misalignment of chromosomes in NudCL2-depleted cells is due to defects in chromosome congression during prometaphase or chromosome segregation at anaphase, we examined the expression and localization of cyclin B1 and the spindle checkpoint proteins Mad2 and Bub1 in these cells. Approximately 90% of NudCL2-depleted mitotic cells with misaligned chromosomes were positive for cyclin B1 throughout the cell and were also positive for Mad2 and Bub1 at kinetochores (Fig 1E-M). Furthermore, live-cell imaging analysis of HeLa cells stably expressing GFP-H2B (GFP-fused histone H2B) revealed that chromosomes failed to align at the metaphase plate in cells depleted of NudCL2 (Fig 1N, Appendix Movie 1 and 2). These results strongly indicate that NudCL2 depletion causes defects in chromosome congression during prometaphase, resulting in the misalignment of chromosomes.

### NudCL2 depletion causes premature sister chromatid separation

To explore how depletion of NudCL2 induces defects in chromosome congression, we first immunostained the centromeric and outer kinetochore markers (CREST and Hec1) on misaligned chromosomes in HeLa cells and found that chromosomes that failed to align at the metaphase plate had one CREST dot associated with only one green Hec1 dot at kinetochores, which is indicative of the loss of sister chromatid cohesion (Fig 2A). To further investigate the function of NudCL2 in sister chromatid cohesion, we prepared mitotic chromosome spreads in HeLa cells transfected with siRNAs for NudCL2, Scc1 or Plk1. Depletion of the cohesin subunit Scc1 induced cohesion loss as previously described [34, 35], and downregulation of Plk1 stabilized cohesion along chromosome arms [7] (Fig 2B-D). A higher frequency of mitotic cells displayed separated sister chromatids in NudCL2-depleted cells compared to that in the control cells, which resembled the Scc1 depletion phenotypes. Similar results were also observed in cells depleted of NudCL2 using another siRNA oligo (Appendix Fig S4) and in HEK-293 cells (Appendix Fig S5). Furthermore, these defects were rescued by ectopic expression of siRNA-resistant NudCL2 in HeLa cells depleted of endogenous NudCL2 (Fig 2E and F). Chromosome spreads followed by immunostaining with CREST and SMC2 (a chromosome arm marker) confirmed that NudCL2 depletion led to precocious sister chromatid separation (Fig 2G and H). Together, these data indicate that NudCL2 plays an essential role in sister chromatid cohesion.

**Figure 2.**
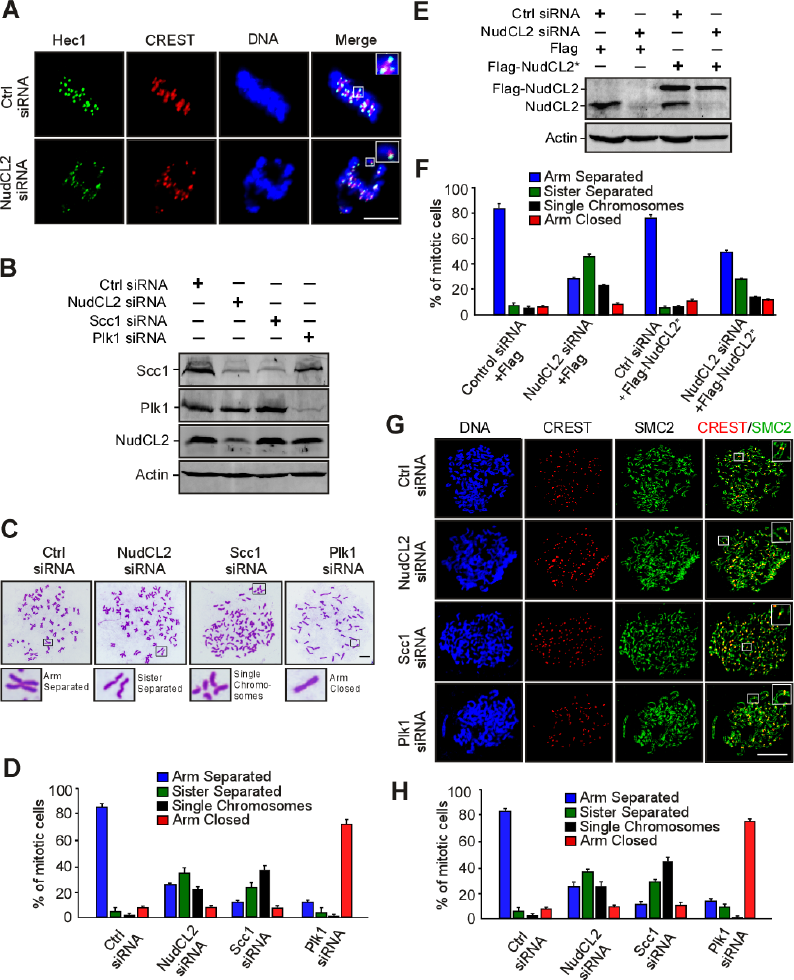
NudCL2 depletion results in premature sister chromatid separation. HeLa cells transfected with the indicated siRNAs and vectors for 72 h were processed for the following analyses. A Immunofluorescence analysis showed that NudCL2-depleted cells exhibited misaligned chromosomes with one CREST dot associated with only one green Hec1 dot at kinetochores. Cells were stained with an anti-Hec1 antibody and human CREST serum. DNA was visualized by DAPI. B-F Chromosome spread assays with Giemsa staining revealed precocious sister chromatid separation in cells depleted of NudCL2. Western blot analysis displayed efficient suppression of NudCL2, Scc1 and Plk1 (B). Actin was used as a loading control. The cells were treated with colcemid for 2.5 h and subjected to chromosome spread analysis (C and D). Ectopic expression of RNAi-resistant NudCL2 (NudCL2*) effectively reversed the defects in sister chromatid separation induced by NudCL2 depletion (E and F). G, H Chromosome spread assay followed by immunofluorescence analysis with the indicated antibodies confirmed the precocious separation of sister chromatids in NudCL2-depleted cells. The cells grown on coverslips were treated with colcemid and subjected to chromosome spread and immunofluorescence analysis. Data information: Mitotic cells in chromosome spread experiments were categorized into four groups according to their chromosomal morphology: (i) arm separated, where the arms of sister chromatids were separated but connected at centromeres; (ii) sister separated (separated sister chromatids), where sister chromatids were separated along the whole chromosome, but their pairing was maintained; (iii) single chromosomes, where sister chromatids were completely separated and scattered; (iv) arm closed, where sister chromatids were connected at both centromeres and arms. Mitotic cells with different chromosomal morphologies were calculated. DNA was visualized with DAPI. Scale bars, 10 μm. Quantitative data are presented as the mean ± SD (at least three independent experiments). More than 150 cells were measured in each experiment. Bars, 10 μm. Higher magnifications of the boxed regions are displayed.

Given that LIS1 is crucial for mitosis and stabilized by NudCL2 [26, 36], we examined whether the role of NudCL2 in sister chromatid cohesion is mediated by LIS1. Chromosome spreads with Giemsa staining clearly revealed that knockdown of LIS1 had no significant effect on sister chromatid cohesion (Appendix Fig S6), suggesting that the defects in sister chromatid cohesion induced by NudCL2 depletion are not due to LIS1 downregulation.

### Depletion of NudCL2 destabilizes cohesin subunits

Since our previous data showed that NudCL2 is involved in the regulation of protein stability [26], and Scc1 protein levels were decreased in NudCL2-depleted cells (Fig 2B), we next explored whether NudCL2 plays a role in the protein stability of cohesin subunits. Depletion of NudCL2 decreased the protein levels of four cohesin subunits, SMC1, SMC3, Scc1 and SA2 (Fig 3A and B). These levels were restored by ectopic expression of siRNA-resistant NudCL2 (Fig 3C). RT-PCR analysis showed that knockdown of NudCL2 had no effect on the mRNA levels of these subunits (Fig 3D). It has been reported that downregulation of other cohesin regulatory factors also induces the precocious separation of sister chromatids in mitosis [3, 37]. We examined the protein levels of two crucial cohesin regulatory factors, Scc4 and Sgo1, in cells depleted of NudCL2 and found no changes (Appendix Fig S7). Thus, these data strongly indicate that NudCL2 plays an essential role in cohesin subunit stability.

**Figure 3.**
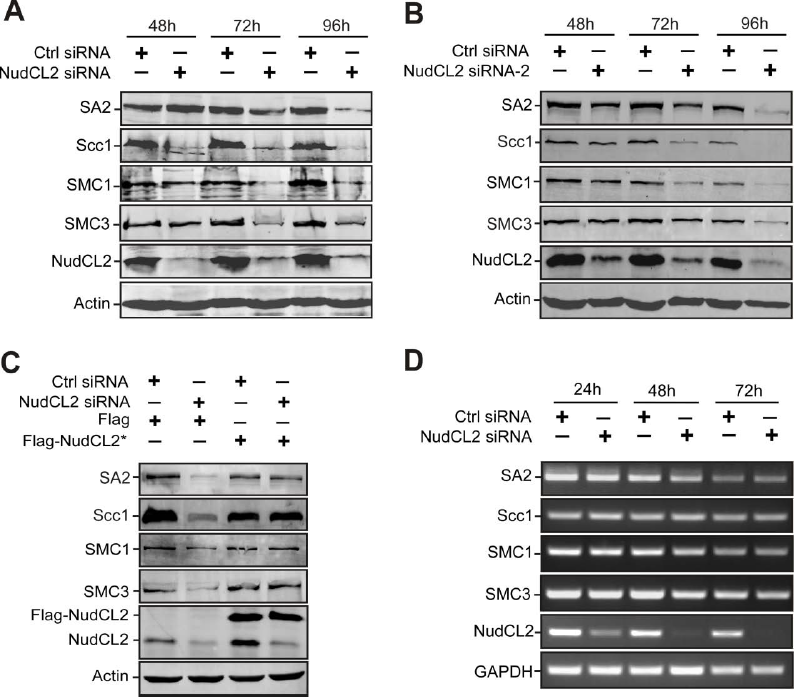
Knockdown of NudCL2 leads to the degradation of cohesin subunits. HeLa cells transfected with the indicated siRNAs and plasmids were subjected to the following analyses. A, B Western blot analysis with the indicated antibodies revealed that NudCL2 depletion by two different NudCL2 siRNAs effectively decreased the protein levels of cohesin subunits. Actin was used as a loading control. C RNAi rescue experiments showed that ectopic expression of NudCL2 rescued the decreased protein levels of cohesin subunits. D Semiquantitative RT-PCR demonstrated that knockdown of NudCL2 had no obvious effect on the mRNA levels of cohesin subunits. GAPDH was used as an internal control.

### Inhibition of Hsp90 leads to precocious sister chromatid separation

Given that NudCL2 facilitates cohesin subunit stability and has a p23 domain that regulates the ATPase activity and chaperoning function of Hsp90 [26, 33], we tried to assess whether Hsp90 is involved in cohesin subunit stability and sister chromatid cohesion in HeLa cells. Our data revealed that inhibition of Hsp90 ATPase activity by geldanamycin (GA) not only obviously decreased the protein levels of cohesin subunits (Fig 4A), but also led to a significant increase in the number of mitotic cells with chromosome misalignment that were positive for cyclin B1 staining (Fig 4B-G). Video microscopy confirmed that inhibition of Hsp90 ATPase activity impaired the alignment of chromosomes at the metaphase plate (Fig 4H, Appendix Movies 3 and 4). Further data showed that inhibition of Hsp90 caused premature sister chromatid separation (Fig 4I and J). Similar results were also observed in HEK-293 cells treated with GA (Appendix Fig S8). Together, these data strongly suggest that Hsp90 is critical for the stability of cohesin subunits and sister chromatid cohesion.

**Figure 4.**
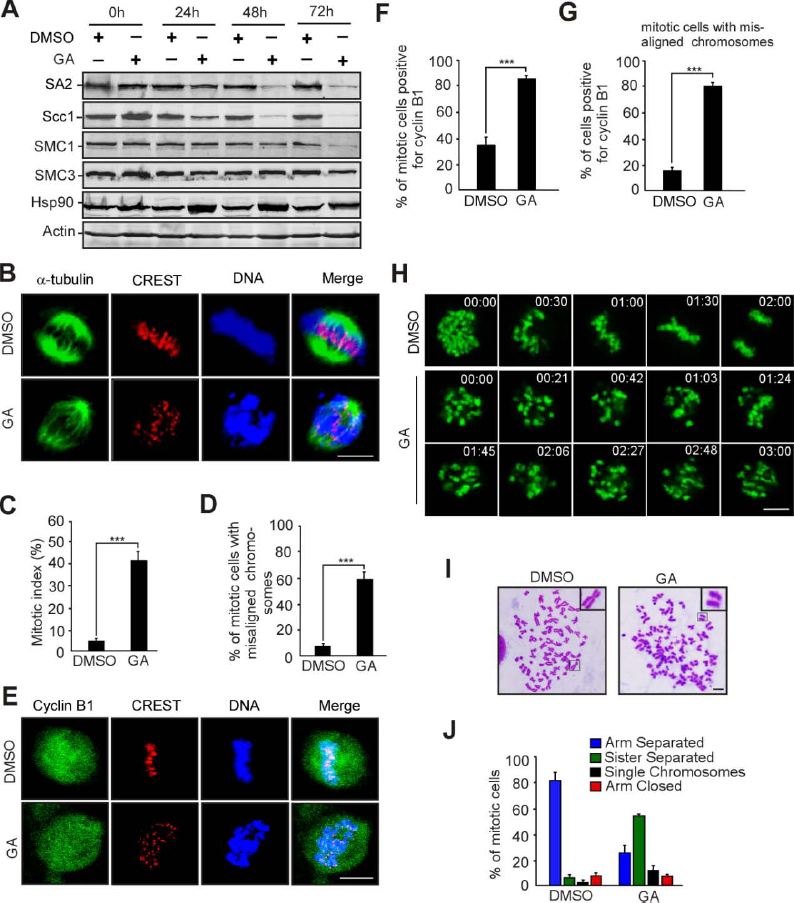
Inhibition of Hsp90 induces defects in sister chromatid cohesion. HeLa cells treated with geldanamycin (GA) or DMSO for 48 h were used for the following analyses. A Western analysis with the indicated antibodies revealed that inhibition of Hsp90 by GA treatment obviously reduced the protein levels of cohesin subunits. Actin, a loading control. B-D Immunofluorescence showed the misalignment of chromosomes in cells treated with GA. The cells were stained with an anti-α-tubulin antibody and CREST serum (B). The mitotic index was calculated (C). The percentage of mitotic cells with misaligned chromosomes was plotted (D). E-G Immunostaining displayed that the majority of GA-treated cells with misaligned chromosomes were positive for cyclin B1. Cells were probed with the indicated antibodies (E). Cyclin B1-positive mitotic cells were quantified (F). The frequency of cyclin B1-positive mitotic cells with misaligned chromosomes was plotted (G). H Live imaging of HeLa cells stably expressing GFP-H2B showed that chromosomes failed to congress to the metaphase plate in GA-treated cells. I, J Cells treated with GA exhibited premature sister chromatid separation. HeLa cells were subjected to chromosome spreads followed by Giemsa staining after treatment with colcemid for 2.5 h (I). Insets, high magnifications of the boxed areas. Mitotic cells with different chromosomal morphologies were counted by the method described in Figure 2 (J). Data information: DNA was visualized with DAPI. Scale bars, 10 μm. Quantitative data are expressed as the mean ± SD (at least three independent experiments). More than 180 cells were scored in each experiment. \*\*\**p* < 0.001, Student’s *t* test.

### NudCL2 regulates cohesin subunit stability via Hsp90

Given that depletion of NudCL2 or inhibition of Hsp90 ATPase activity destabilized cohesin subunits and caused precocious sister chromatid separation (Figs 2–4), we speculated that NudCL2 may be involved in the regulation of cohesin stability via Hsp90. We designed a series of rescue experiments and found that the exogenous expression of Hsp90 efficiently reversed the instability of cohesin subunits and the defects in sister chromatid cohesion induced by NudCL2 depletion (Fig 5A and B). In contrast, ectopic expression of NudCL2 failed to reverse the degradation of cohesin subunits and the defects in sister chromatid cohesion caused by Hsp90 inhibition (Fig 5C and D). Furthermore, depletion of NudCL2 had no synergistic effect with Hsp90 inhibition on cohesin subunits stability or sister chromatid cohesion (Fig 5E and F). Thus, these data imply that NudCL2 modulates the stability of cohesin subunits via Hsp90.

**Figure 5.**
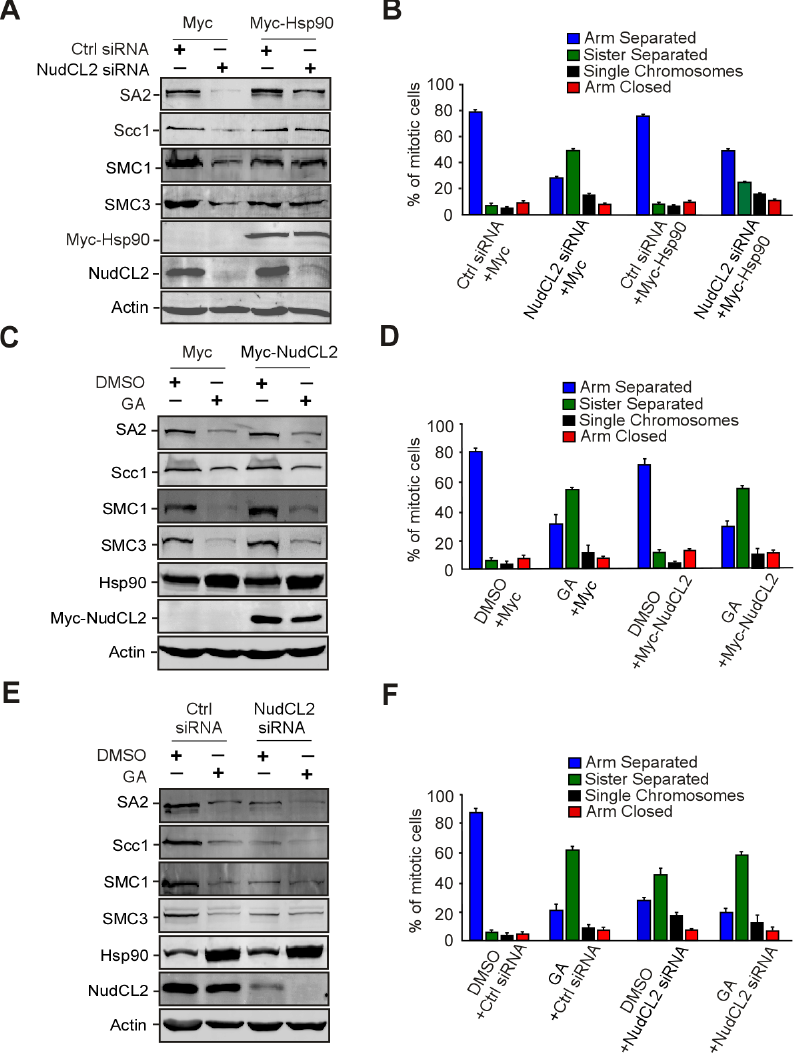
**Ectopic expression of Hsp90 rescues the defects caused by NudCL2 depletion but not** ***vice versa.*** HeLa cells transfected with the indicated siRNAs and vectors were treated with geldanamycin and subjected to Western blotting and chromosome spreads followed by Giemsa staining. Actin was used as a loading control. The percentages of mitotic cells with different chromosomal morphologies were measured by the method described in Figure 2. Quantitative data are shown as the mean ± SD (at least three independent experiments). More than 170 cells were calculated in each experiment. A, B Exogenous expression of Hsp90 reversed the degradation of cohesin subunits and premature sister chromatid segregation in NudCL2-depleted cells. C, D Ectopic expression of NudCL2 was unable to rescue the decreased levels of cohesin subunits and defects in sister chromatid cohesion in cells with Hsp90 inhibition. E, F Depletion of NudCL2 had no synergistic effects with Hsp90 inhibition on cohesin subunit stability and sister chromatid cohesion.

### NudCL2 interacts with Hsp90 and cohesin subunits

To investigate how NudCL2 regulates cohesin subunits stability by Hsp90, we first examined the potential interaction among NudCL2, Hsp90 and cohesin subunits. GST pull-down assays using cell lysates revealed that GST-NudCL2 bound to Hsp90 and cohesin subunits (Fig 6A). Further immunoprecipitation experiments with anti-NudCL2 or Scc1 antibody showed the association of endogenous NudCL2, Hsp90 and cohesin subunits *in vivo* (Fig 6B and C). To determine whether NudCL2 directly interacted with cohesin subunits, we purified recombinant cohesin subunits (His-SMC1, His-SMC3, His-Scc1 and His-SA2) from insect Sf9 cells using Bac-to-Bac baculovirus expression system. The data showed that GST-NudCL2 protein directly interacted with SMC1 and SMC3, but not Scc1 and SA2 *in vitro* (Fig 6D-G).

**Figure 6.**
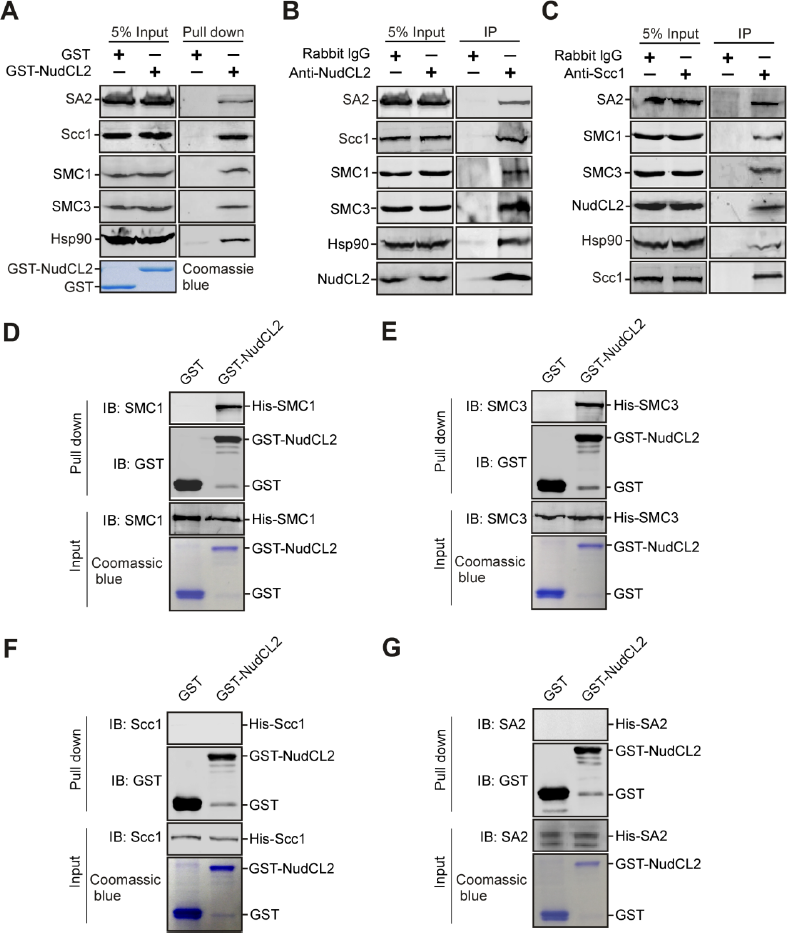
**NudCL2 interacts with cohesin subunits and Hsp90** ***in vitro*** **and** ***in vivo.*** A GST pull-down assays showed that NudCL2 associated with cohesin subunits and Hsp90 *in vitro.* Purified GST or GST-NudCL2 protein was incubated with HeLa cell lysates and subjected to immunoblotting. The inputs of GST and GST-NudCL2 were stained with Coomassie brilliant blue. B, C Endogenous NudCL2 bound to four cohesin subunits and Hsp90 *in vivo.* Total lysates of HeLa cells were immunoprecipitated with the indicated antibodies or IgGs and processed for Western blotting. D-G NudCL2 protein directly interacted with SMC1 and SMC3 but not Scc1 or SA2 *in vitro.* His-tagged Scc1, SA2, SMC1 and SMC3 were expressed in Sf9 insect cells and purified with nickel-nitrilotriacetic acid beads. GST or GST-NudCL2 protein was incubated with each of purified cohesin subunits and subjected to Western analysis with the indicated antibodies.

### NudCL2 functions as an Hsp90 cochaperone

Since NudCL2 not only regulates the stability of cohesin subunits and LIS1, but also contains a conserved p23 domain, the core structure of p23 required for its cochaperone function of Hsp90 [26, 33], we explored whether NudCL2 functions as an Hsp90 cochaperone to stabilize client proteins. We first purified NudCL2, p23 and Hsp90 proteins, and performed *in vitro* Hsp90 ATPase assays (Fig 7A). The results revealed that both NudCL2 and p23 significantly inhibited the ATPase activity of yeast Hsp90 (Fig 7B). Further heat-induced aggregation experiments with two Hsp90 substrates, citrate synthase (CS) and luciferase, showed that NudCL2 enhanced the chaperone function of Hsp90 to inhibit the aggregation of CS and luciferase, whereas NudCL2 itself did not suppress their heat-induced aggregation (Fig 7C and D).

**Figure 7.**
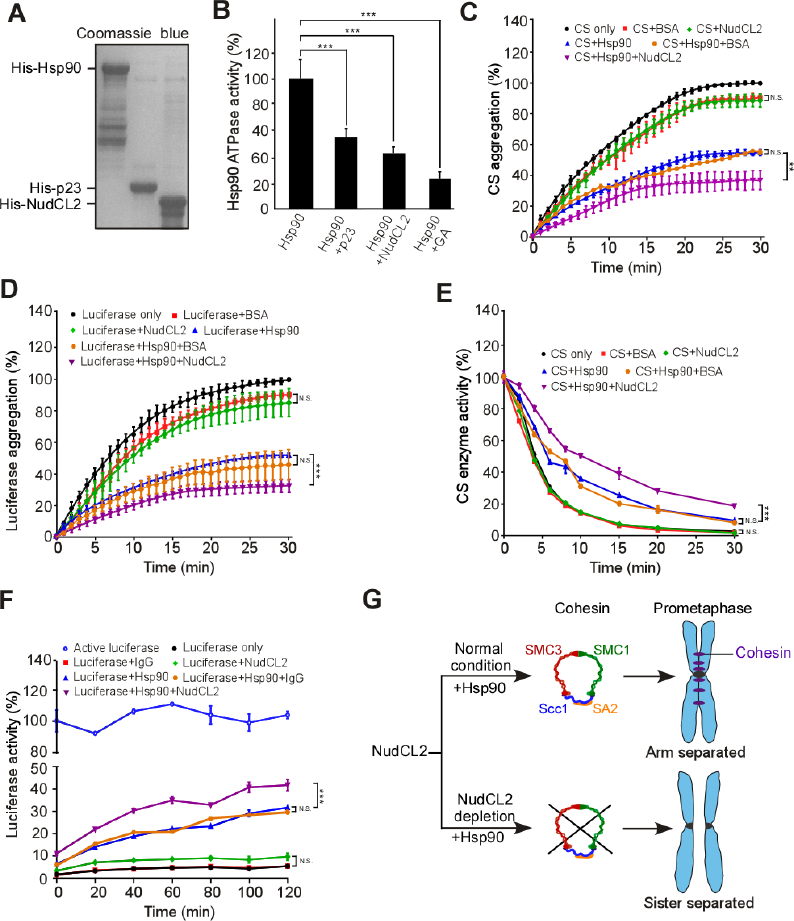
NudCL2 enhances the chaperone activity of Hsp90. A Purified His-tagged p23, Hsp90, and NudCL2 proteins were subjected to SDS-PAGE with Coomassie blue staining. B NudCL2 inhibited the *in vitro* ATPase activity of Hsp90. Hsp90 ATPase activity was measured in the presence of p23, NudCL2 or geldanamycin and presented as the activity relative to that of only Hsp90. C, D NudCL2 suppressed the heat-induced aggregation of citrate synthase (CS) and luciferase with Hsp90. CS (C) and luciferase (D) were incubated with the indicated proteins for different times. Aggregation was determined by light scattering (370 nm) at each time point and calculated as the percentage of CS or luciferase aggregation after a 30 min incubation. E NudCL2 reduced the rate of heat-induced CS inactivation with Hsp90. CS was incubated with the indicated proteins at 43ºC for different times. The enzymatic activity of CS was detected at 30ºC by light scattering at 412 nm. All activities were expressed as a percentage of the initial activity of CS (0 min). F NudCL2 facilitated the refolding of luciferase after heat-induced aggregation. Luciferase was incubated at 42ºC for 15 min in the presence of the indicated proteins. The enzymatic activity of luciferase was monitored by a luminometer at each time point. The activity was calculated as a percentage of the activity of luciferase at 22 ºC after 15 min incubation. G Working model for the role of NudCL2 in sister chromatid cohesion. NudCL2 stabilizes cohesin subunits by acting as an Hsp90 cochaperone to enhance cohesin stability. Depletion of NudCL2 induces cohesin subunits degradation and results in premature sister chromatid segregation during mitosis. Data information: Quantitative data are presented as the mean ± SD (at least three independent experiments). \*\*\**p* < 0.001, Student’s *t* test.

Both CS and luciferase enzymes are active in their native state and lose activity upon unfolding or denaturation, and this property has been utilized to study the chaperoning function of Hsp90 influenced by its cochaperones [38, 39]. To confirm whether NudCL2 acts as an Hsp90 cochaperone to regulate the enzymatic activity of Hsp90 client proteins, we carried out CS thermal inactivation and luciferase refolding assays *in vitro*. The results showed that upon Hsp90, NudCL2 not only significantly reduced the heat-induced CS inactivation (Fig 7E), but also promoted the luciferase activity recovery after heat-induced inactivation (Fig 7F). Interestingly, in the absence of Hsp90, NudCL2 did not suppress the heat-induced CS inactivation or enhance the recovery of luciferase activity after heat-induced inactivation (Fig 7E and 7F). Taken all together, these data indicate that NudCL2 may function as an Hsp90 cochaperone to enhance the stability and function of client proteins.

## Discussion

The Hsp90 cochaperone p23 plays a critical role in the stabilization of client proteins, including steroid hormone receptors and telomerase, by inhibiting Hsp90 ATPase activity [33, 40, 41]. In this report, we provide evidence that NudCL2 interacts with Hsp90 and suppresses the ATPase activity of Hsp90 (Fig 6A-6C and Fig7A and 7B). Further results reveal that NudCL2 inhibits the heat-induced aggregation and enzymatic inactivation of Hsp90 substrates (Fig 7C-F). Moreover, we also show that either NudCL2 depletion or Hsp90 inhibition decreases the stability of cohesin subunits and induces premature sister chromatid separation (Figs 2–4). Ectopic expression of Hsp90 efficiently rescues the defects induced by NudCL2 depletion, but not *vice versa* (Fig 5). These results indicate that the p23 domain-containing protein NudCL2 is an Hsp90 cochaperone to stabilize cohesin subunits (Fig 7G).

*NudC* was first identified in *Aspergillus nidulans* as a key upstream regulator of the LIS1/dynein complex [20, 23, 24]. Our group and others have found that vertebrate NudC has three principal homologs, NudC, NudCL and NudCL2, all of which contain the conserved p23 domain [25, 26, 28, 31, 32]. NudC is not only able to interact with Hsp90 and regulate LIS1 stability via the Hsp90 pathway, but also stabilizes cofilin 1 to regulate ciliogenesis in an Hsp90-independent manner [28, 32]. Additionally, NudCL has been documented to play an important role in the stability regulation of dynein intermediate chain [25]. Recently, our data show that NudCL2 promotes LIS1 stability by enhancing the interaction between Hsp90 and LIS1 [26]. In this report, we further find that NudCL2 acts as an Hsp90 cochaperone to stabilize cohesin subunits by modulating the ATPase activity of Hsp90. Intriguingly, NudC and NudCL, but not NudCL2, possess an intrinsic chaperoning activity independent of Hsp90 to regulate the stability of client proteins [25, 31, 32]. Taken together, these data imply that the members of NudC family may act as a new type of regulator of protein stability possibly via the Hsp90-dependent or -independent pathway.

Emerging studies have demonstrated the molecular regulation underlying the stability of the cohesin-DNA interaction [4–7, 16]. However, how the stability of cohesin subunits is regulated remains unclear. Recent studies showed that SMC3 knockdown leads to SMC1 instability, but SMC1 depletion has no significant effect on SMC3 stability [42]. Downregulation of either SMC1 or SMC3 causes Scc1 degradation, while depletion of Scc1 only decreases SA protein levels [42, 43]. Here, our data reveal that knockdown of either SMC3 or SMC1 reduces the stability of the other three cohesin subunits (Appendix Fig S9). Scc1 depletion decreases the protein levels of SA2, but not SMC1 or SMC3. SA2 depletion does not affect the stability of the other three subunits. These results indicate that either SMC1 or SMC3 is essential for the stability of Scc1 and SA2, but not *vice versa*. Moreover, we find that depletion of NudCL2 decreases the protein levels of all cohesin subunits (Fig 3). Purified GST-NudCL2 directly interacts with His-tagged SMC1 and SMC3, but not Scc1 or SA2 (Fig 6D-G). Together, these data suggest that NudCL2 may promote the stability of cohesin subunits by directly interacting with SMC1 and SMC3.

In summary, this study provide evidence that NudCL2 is essential for sister chromatid cohesion by acting as an Hsp90 cochaperone to stabilize cohesin subunits. Depletion of NudCL2 causes cohesin subunits degradation and results in premature sister chromatid segregation during mitosis. This work reveals a previously uncharacterized mechanism for the regulation of chromosome segregation in mammalian cells.

## Materials and Methods

### Plasmids and small interfering RNAs (siRNAs)

Human Flag-NudCL2, Flag-NudCL2* (with a silent mutation of three nucleic acids in the RNAi targeting region: ACCTTGA<u>A</u>AA<u>G</u>T<u>G</u>ACTGCT), GST-NudCL2, His-p23 and yeast His-Hsp90α vectors were constructed previously [26, 28]. Human Hsp90α was cloned by RT-PCR and inserted into pcDNA 3.1/Myc-His C (Myc/His-tag vector, Invitrogen). Full-length NudCL2 was cloned by PCR using Flag-NudCL2 as a template and subcloned into pET-28a (His-tag vector, Novagen) and pcDNA 3.1/Myc-His C. Full-length human SMC1, SMC3, Scc1 and SA2 cloned by RT-PCR were inserted into pFastBac-HT A (His-tag vector, Invitrogen). All of these constructs were confirmed by DNA sequencing.

All siRNAs were synthesized by Genepharma. The sequences of the sense strands of the siRNA duplexes are as follows:

LIS1: 5’-GAACAAGCGAUGCAUGAAGTT-3’ [36];

NudCL2: 5’-ACCUUGAGAAAUAACTGCUTT-3’;

NudCL2-2: 5’-CAAGGGCAAACUCUUUGAUTT-3’;

Plk1: 5’-GGGCGGCUUUGCCAAGUGCTT-3’ [8];

SA2: 5’-GUACGGCAAUGUCAAUAUATT-3’ [44];

Scc1: 5’-AUACCUUCUUGCAGACUGUTT-3’ [8];

SMC1: 5’- GGAAGAAAGUAGAGACAGATT-3’ [45];

SMC3: 5’-UGUCAUGUUUGUACUGAUATT-3’ [46].

### Cell culture and transfections

HeLa and HEK-293 cells were maintained in Dulbecco’s Modified Eagle’s Medium (DMEM, Corning) supplemented with 10% fetal bovine serum (Invitrogen) at 37ºC in 5% CO_2_. Plasmids were transfected with PolyJet (SignaGen), and the siRNA duplexes were transfected with Lipofectamine RNAiMAX (Invitrogen). The transfection processes were performed according to the manufacturer’s instructions.

### Drug treatments

Geldanamycin (GA, Tocris) was stored at −20°C as a stock solution of 1.78 mM in DMSO. Cells were treated with geldanamycin for the indicated times as described in the text. The final concentration of GA was 1.78 μM for HeLa cells and 0.5 μM for HEK-293 cells. For chromosome spread experiments, 100 ng/ml of colcemid (Sigma) was used to treat cells for 2.5 h.

### Antibodies

For immunofluorescence, antibodies against α-tubulin (Sigma, T6199), cyclin B1 (Santa Cruz, sc-70898), Bub1 (Abcam, ab900), Mad2 (Santa Cruz, sc-28261), Hec1 (Abcam, ab3613) and SMC2 (Abcam, ab10412) and human CREST autoimmune serum (Antibodies, Inc., 15-235-0001) were used. For Western blot analysis, antibodies against Scc1 (Bethyl, A300-080A), SA2 (Bethyl, A300-581A), SMC1 (Bethyl, A300-055A), SMC3 (Bethyl, A300-060A), Scc4 (Abcam, ab46906), Sgo1 (Santa Cruz, sc-54329), Plk1 (Santa Cruz, sc-53418) and Hsp90 (Abcam, ab59459) were utilized. The anti-NudCL2 antibody was generated as described previously [19]. An antibody against actin (Sigma, T1978) was used as the loading control. The secondary antibodies for immunofluorescence analyses were Alexa Fluor 488-conjugated anti-rabbit or anti-mouse IgG and Alexa Fluor 568-conjugated anti-human IgG (Invitrogen). Goat anti-mouse or anti-rabbit secondary antibodies (LI-COR) conjugated with either Alexa Fluor 680 or IRDye 800 was used for Western blot analysis.

### Immunofluorescence staining

HeLa or HEK-293 cells grown on glass coverslips were fixed for 15 min with cold methanol (−20°C) and then incubated with primary antibodies for 2 h and secondary antibodies for 1 h at room temperature. DNA was stained with DAPI (Sigma). The mounted coverslips were analyzed by confocal fluorescence microscopy with an oil immersion 60× objective (LSM510, Zeiss).

### Live-cell images

HeLa cells stably expressing GFP-H2B were treated with either siRNA for 72 h or GA for 48 h. Then, the cells were analyzed with time-lapse video microscopy to track the cell cycle progression. Fluorescence images were captured at 3-min intervals for at least 3 h with LSM510 software (Zeiss). The videos were further edited by ImageJ software (NIH).

### GST pull-down assays

GST pull-down assays were performed as described previously [27, 47]. GST and GST-NudCL2 were purified from bacteria. His-tagged SMC1, SMC3, Scc1 and SA2 were purified from insect Sf9 cells using the Bac-to-Bac baculovirus expression system (Invitrogen). To detect the association among NudCL2, cohesin subunits and Hsp90, the blots were probed with the respective antibodies as indicated in the text.

### Immunoprecipitation and Western blot assays

Immunoprecipitation for endogenous proteins was performed as described previously [27]. Briefly, whole-cell extracts were generated in TBSN buffer (20 mM Tris [pH 8.0], 150 mM NaCl, 0.5% Nonidet P-40, 5 mM EGTA, 1.5 mM EDTA, 0.5 mM Na_3_VO_4_, 20 mM p-nitrophenyl phosphate) supplemented with protease inhibitors and subjected to coimmunoprecipitation analysis with the indicated antibodies. Western blot analyses were performed with the antibodies as indicated and analyzed using the LI-COR Odyssey (LI-COR) system.

### Chromosome spread assays

Chromosome spread assays were prepared as previously described [48]. In brief, cells were collected and then treated with hypotonic solution (DMEM: H_2_O at a ratio of 2:3) for 5.5 min at room temperature and fixed with freshly prepared Carnoy’s solution (75% methanol, 25% acetic acid). Cells in Carnoy’s solution were dropped onto glass slides, stained with 5% Giemsa (Solarbio) and analyzed by bright field microscopy (BX51, Olympus). To perform chromosome spreads for immunofluorescence staining, HeLa cells grown on coverslips were treated with 75 mM KCl hypotonic solution for 15 min at 37°C, and centrifuged for 5 min at 2,000 g to spread chromosomes onto coverslips. The samples on coverslips were washed with PHEM buffer (60 mM PIPES, 25 mM HEPES, 10 mM EGTA, 2 mM MgCl_2_, pH 6.9), fixed for 5 min with cold methanol (−20°C) and processed for immunofluorescence microscopy (LSM510, Zeiss).

### Hsp90 ATPase assays

Hsp90 ATPase assays were performed as described previously [28]. Briefly, His-Hsp90 (1 µM) was incubated with His-NudCL2 (1 µM), His-p23 (1 µM) or geldanamycin (1.78 µM) at 37°C for 20 min in the reaction buffer (50 mM Tris, pH 7.4, 20 mM KCl, 6 mM MgCl_2_, 1 mM dithiothreitol, 0.5 mM ATP). The released inorganic phosphate was determined by measuring the absorbance at 650 nm using the Cyto Phosphate Assay BIOCHEM kit (Cytoskeleton).

### Aggregation assays

The aggregation reactions of CS (Sigma) or luciferase (Promega) were carried out as described previously [49]. Briefly, CS (0.15 μM) or luciferase (0.15 μM) was incubated alone or with BSA (0.15 μM), NudCL2 (0.15 μM) or Hsp90 (0.15 μM) at 43°C (for CS) or 42°C (for luciferase) for 30 min in 40 mM HEPES-KOH (pH 7.5). To monitor the kinetics of thermal aggregation, light scattering was measured at 370 nm by a DU 800 spectrophotometer (Beckman).

### CS thermal inactivation assays

The enzyme inactivation assay of CS was performed as described previously [49, 50]. Briefly, CS (0.15 μM) was incubated at 43°C in the absence or presence of IgG (1.2 μM), NudCL2 (0.6 μM) or Hsp90 (0.6 μM) in the inactivation buffer (40 mM HEPES-KOH, 0.1mM EDTA, pH 7.5). Aliquots (100 μl) was taken at the indicated times and mixed with 650 μl of 100 mM Tris (pH 8.1), 50 μl of 3 mM acetyl-CoA (Sigma), 100 μl of 1 mM DTNB (Sigma), and 100 μl of 5 mM oxaloacetate (Sigma), and then incubated at 30°C for 1 min to eliminate the false readings. To monitor the CS activity, the readings were measured at 30°C for 1 min with 20-s intervals at 412 nm by SpectraMax (Molecular Devices).

### Luciferase refolding assays

Luciferase refolding assays were carried out as described previously [49]. Luciferase (0.2 μM) was incubated either alone or in the presence of IgG (2 μM), NudCL2 (0.2 μM) or Hsp90 (0.2 μM) at 22°C or 42°C for 15 min in the refolding buffer (5 mM MgCl_2_, 10 mM KCl, 2 mM DTT, 50 mM HEPES-KOH, pH 7.5). After incubation, 10 μl of each mixture was immediately added to a solution containing 18 μl of rabbit reticulocyte lysate (RRL) (Promega) and 2 μl of 0.1 M ATP. During the incubation at 30°C, the luciferase enzyme activity was measured by SpectraMax (Molecular Devices) using the Luciferase Assay System (Promega) at the indicated times. The tubes were treated with 1 mg/ml BSA for 15 min to prevent luciferase adsorption to the walls.

### Statistical analysis

Data are representative of at least three independent experiments. The mean and standard deviations (SD) were calculated for all quantitative experiments. Student’s t-test was used to determine statistically significant differences between groups.

#### Acknowledgements

The authors are grateful to Dangsheng Li and Xueliang Zhu for helpful suggestions, Lynn W. Enquist for his comments on the manuscript. We thank the members of Zhou’s lab for the helpful comments and suggestions during the work. This work was supported by the National Natural Scientific Foundation of China (31671394, 31471259, 31701214, 91740205, 31620103911, 31571446 and 31771540) and the National Key Research and Development Program of China (2016YFA0100301).

### Author contributions

YY, WW, and ML designed and performed the majority of the experiments. WZ provided technical assistance for the aggregation assays. YH provided assistance for GST pull-down assays. YG, WZ, XY, WL, and FW discussed the results and commented on the manuscript. YY and WW wrote the original draft. DC and TZ edited the manuscripts. TZ supervised the project.

## Appendix

**Appendix Figure S1.**
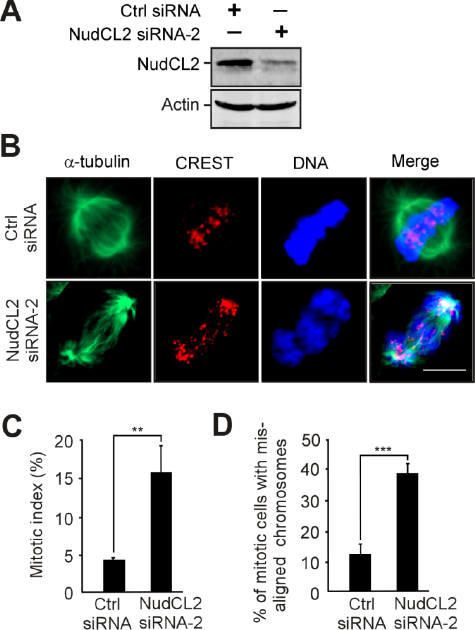
NudCL2 depletion induces mitotic defects in HeLa cells. HeLa cells were transfected with control or NudCL2 siRNA-2 for 72 h and processed for the following experiments. A Western analysis showed depletion of NudCL2. Actin was used as a loading control. B-D Immunofluorescence of NudCL2-depleted cells exhibited chromosome misalignment. Cells were stained with the indicated antibodies (B). DNA was labeled with DAPI. Scale bars, 10 μm. Mitotic cells were counted and the mitotic index was calculated (C). The percentage of mitotic cells with misaligned chromosomes was higher in NudCL2-depleted cells than in control cells (D). Data information: Quantitative data are expressed as the mean ± SD (at least three independent experiments). More than 200 cells were counted in each experiment. \*\**p* < 0.01; \*\*\**p* < 0.001, Student’s *t* test.

**Appendix Figure S2.**
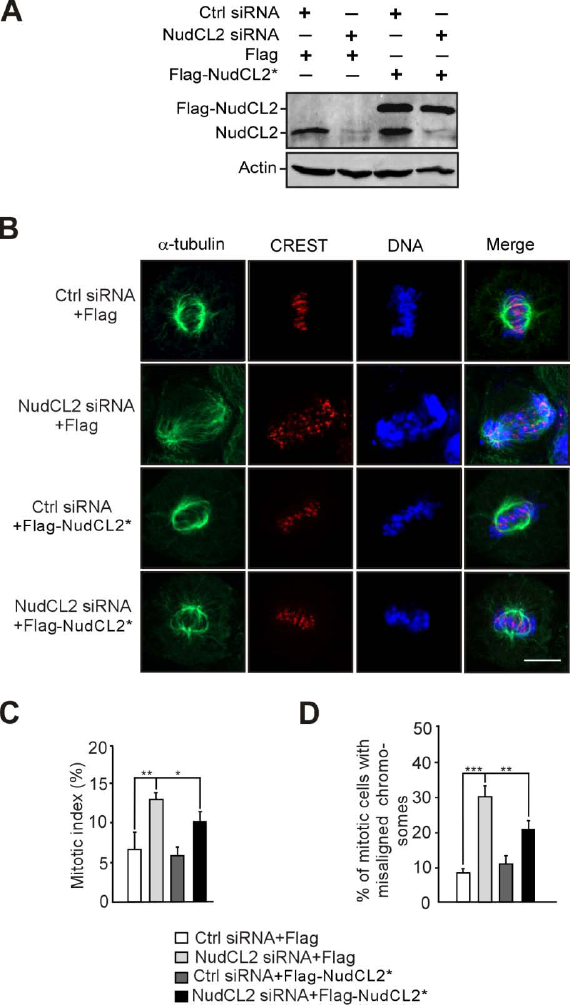
Ectopic expression of RNAi-resistant NudCL2 reverses mitotic defects induced by NudCL2 depletion. HeLa cells transfected with the indicated siRNAs and vectors were subjected to the following analyses. A Western analysis showed efficient suppression of endogenous NudCL2 and ectopic expression of NudCL2. Actin, a loading control. B-D Immunofluorescence revealed that ectopic expression of RNAi-resistant NudCL2 (NudCL2*) significantly reversed the mitotic defects caused by NudCL2 depletion. Cells were stained with the indicated antibodies (B). DNA was labeled with DAPI. Mitotic index was calculated (C). The percentage of mitotic cells with misaligned chromosomes was measured (D). Data information: Quantitative data are expressed as the mean ± SD (at least three independent experiments). More than 150 cells were counted in each experiment. \**p* < 0.05; \*\**p* < 0.01; \*\*\**p* < 0.001, Student’s *t* test.

**Appendix Figure S3.**
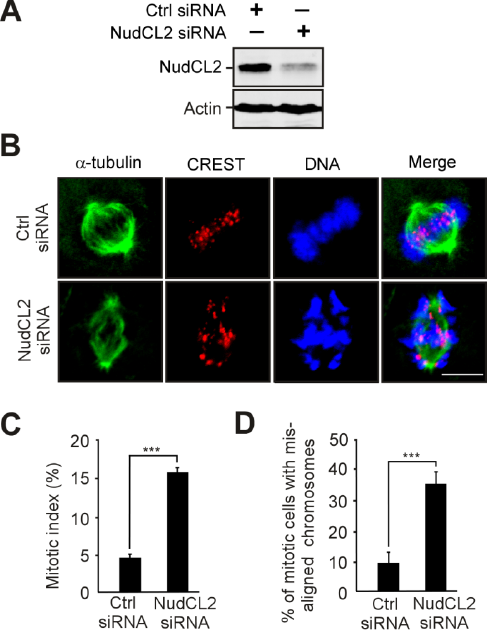
NudCL2 depletion induces mitotic defects in HEK-293 cells. HEK-293 cells were transfected with control or NudCL2 siRNA for 72 h and subjected to the following analyses. A Western blotting showed obvious depletion of NudCL2. Actin was used as a loading control. B-D Immunofluorescence analysis of NudCL2-depleted cells exhibited misalignment of chromosomes. Cells were stained with the indicated antibodies (B). DNA was visualized by DAPI. Mitotic index was calculated (C). The percentage of mitotic cells with misaligned chromosomes was higher in NudCL2-depleted cells than in control cells (D). Data information: Quantitative data are presented as the mean ± SD (at least three independent experiments). More than 200 cells were calculated in each experiment. \*\*\**p* < 0.001, Student’s *t* test.

**Appendix Figure S4.**
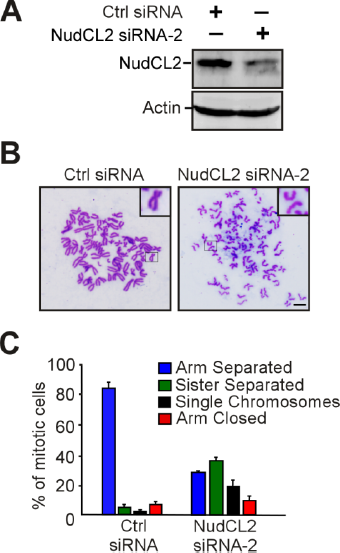
NudCL2 depletion results in premature sister chromatid separation in HeLa cells. HeLa cells were transfected with control or NudCL2 siRNA-2 for 72 h and subjected to the following analyses. A Immunoblotting showed substantial decrease of NudCL2. Actin, a loading control. B, C Depletion of NudCL2 induced precocious sister chromatid separation. The cells were treated with colcemid for 2.5 h and subjected to chromosome spreads and Giemsa staining. Representative images are shown (B). Insets, high magnifications of the boxed areas. The frequencies of four chromosomal morphologies were measured using the method described in Fig 2 (C). Data information: Quantitative data are expressed as the mean ± SD (at least three independent experiments). More than 200 cells were scored in each experiment.

**Appendix Figure S5.**
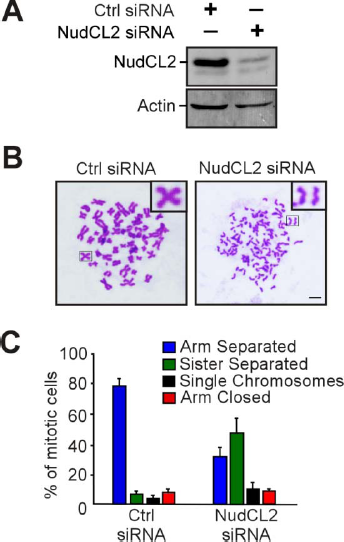
Depletion of NudCL2 promotes premature sister chromatid separation in HEK-293 cells. HEK-293 cells were transfected with control or NudCL2 siRNA for 72 h and subjected to the following analyses. A Immunoblotting displayed efficient suppression of NudCL2. Actin was used as a loading control. B, C Depletion of NudCL2 led to precocious sister chromatid separation. The cells were treated with colcemid and prepared for chromosome spreads followed by Giemsa staining (B). Representative images are shown. Insets, high magnifications of the boxed areas. The percentage of cells with different chromosomal morphologies was measured using the method described in Fig 2 (C). Data information: Quantitative data are expressed as the mean ± SD (at least three independent experiments). More than 200 cells were measured in each experiment.

**Appendix Figure S6.**
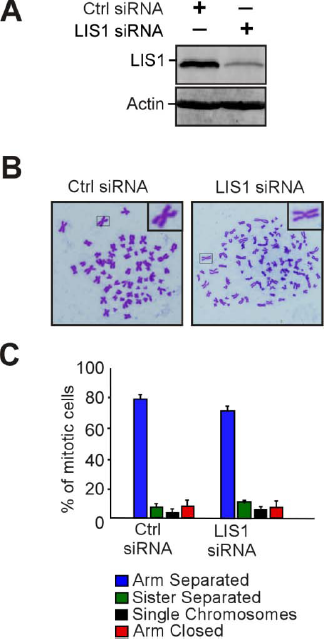
Depletion of LIS1 has no obvious effect on sister chromatid cohesion. HeLa cells were transfected with control or LIS1 siRNA for 72 h and subjected to the following analyses. A Western analysis showed efficient suppression of LIS1. Actin was used as a loading control. B, C Chromosome spreads followed by Giemsa staining revealed that cells depleted of LIS1 displayed no obvious deficiencies in sister chromatid cohesion. Representative images are shown (B). Insets, high magnifications of the boxed areas. The frequencies of four chromosomal morphologies were measured by the method described in Fig 2 (C). Data information: Quantitative data are expressed as the mean ± SD (at least three independent experiments). More than 200 cells were scored in each experiment.

**Appendix Figure S7.**
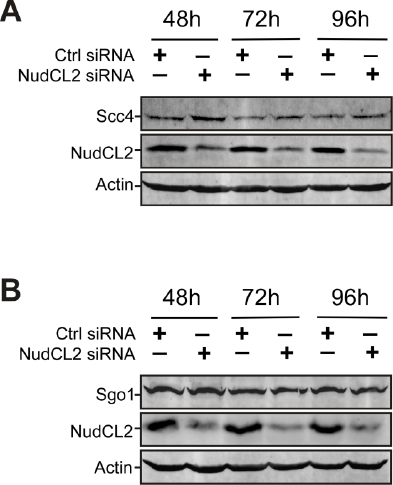
Knockdown of NudCL2 does not affect the protein levels of Scc4 and Sgo1. HeLa cells were transfected with control or NudCL2 siRNA for the indicated times and subjected to Western blotting with the indicated antibodies. Depletion of NudCL2 had no effect on the protein levels of Scc4 (A) and Sgo1 (B). Actin, a loading control.

**Appendix Figure S8.**
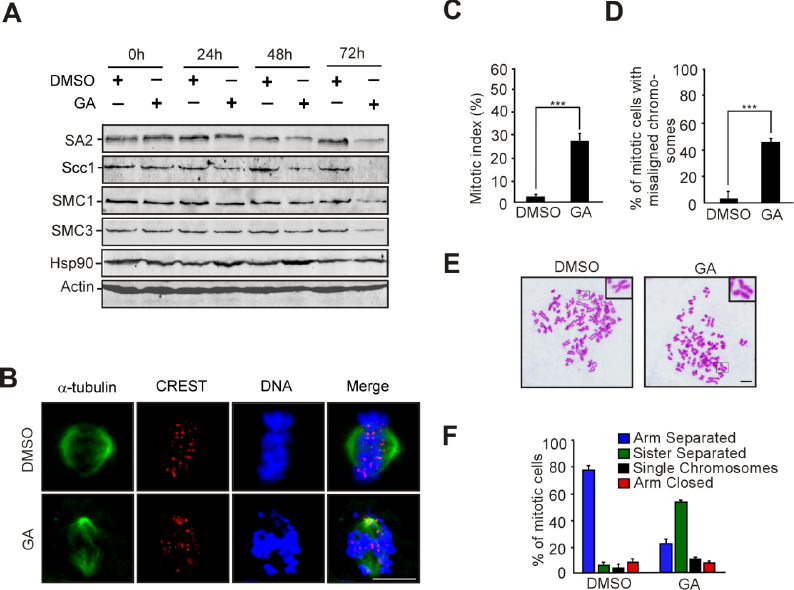
Inhibition of Hsp90 causes defects in sister chromatid cohesion in HEK-293 cells. HEK-293 cells treated with GA or DMSO were subjected to the following analyses. A Inhibition of Hsp90 decreased the levels of cohesin subunits. The cells treated with GA or DMSO were harvested at the indicated times and subjected to immunoblotting analysis with the antibodies as shown. Actin, a loading control. B-D Immunofluorescence showed obvious chromosome misalignment in cells treated with GA for 48 h. Cells were stained with the indicated antibodies (B). Mitotic index was calculated (C). The percentage of mitotic cells with misaligned chromosomes was calculated (D). E, F Inhibition of Hsp90 caused premature sister chromatid separation. The cells were treated with colcemid for 2.5 h and subjected to chromosome spreads and Giemsa staining (E). Representative images are shown. Insets, high magnifications of the boxed areas. The percentages of cells with different chromosomal morphologies were determined as described in Fig 2 (F). Data information: Quantitative data are presented as the mean ± SD (at least three independent experiments). More than 200 cells were scored in each experiment. \*\*\**p* < 0.001, Student’s *t* test.

**Appendix Figure S9.**
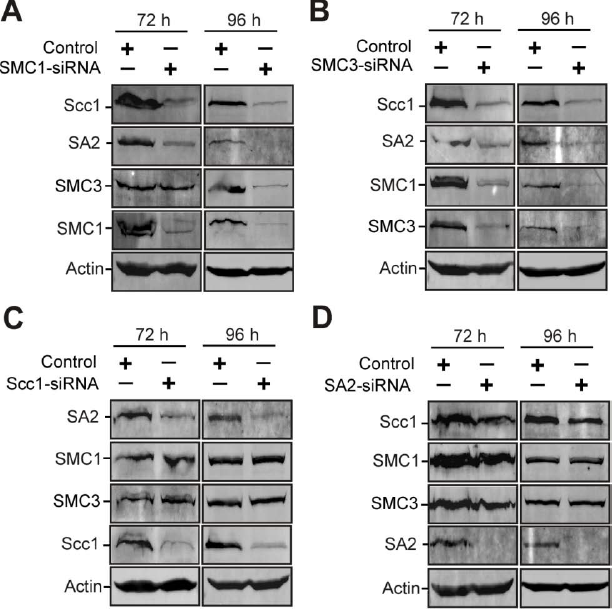
The interdependence of cohesin subunits stability. HeLa cells were transfected with control or the indicated siRNAs and subjected to Western analysis. A SMC1 depletion decreased the stability of SMC3, Scc1 and SA2. B Knockdown of SMC3 reduced the protein levels of SMC1, Scc1 and SA2. C Downregulation of Scc1 caused the decrease in the protein level of SA2, but not SMC1 or SMC3. D Depletion of SA2 had no effect on the other cohesin subunits. Actin, a loading control.

**Appendix Movie S1. Mitotic progression of control cell stably expressing GFP-H2B.**

**Appendix Movie S2. Mitotic progression of NudCL2-depleted cell stably expressing GFP-H2B.**

**Appendix Movie S3. Mitotic progression of DMSO-treated HeLa cell stably expressing GFP-H2B.**

**Appendix Movie S4. Mitotic progression of geldanamycin-treated HeLa cell stably expressing GFP-H2B.**

